# Dynamic Graphical Models of Molecular Kinetics

**DOI:** 10.1101/467050

**Authors:** Simon Olsson, Frank Noé

## Abstract

Most current molecular dynamics simulation and analysis methods rely on the idea that the molecular system can be characterized by a single global state, e.g., a Markov State in a Markov State Model (MSM). In this approach, molecules can be extensively sampled and analyzed when they only possess a few metastable states, such as small to medium-sized proteins. However this approach breaks down in frustrated systems and in large protein assemblies, where the number of global meta-stable states may grow exponentially with the system size. Here, we introduce Dynamic Graphical Models (DGMs), which build upon the idea of Ising models, and describe molecules as assemblies of coupled subsystems. The switching of each sub-system state is only governed by the states of itself and its neighbors. DGMs need many fewer parameters than MSMs or other global-state models, in particular we do not need to observe all global system configurations to estimate them. Therefore, DGMs can predict new, previously unobserved, molecular configurations. Here, we demonstrate that DGMs can faithfully describe molecular thermodynamics and kinetics and predict previously unobserved metastable states for Ising models and protein simulations.

## Introduction

How many states does a macromolecule have? When is a molecular dynamics (MD) simulation converged? State-based MD analysis methods such as Markov State Models (MSMs)^1–5^ and related MD simulation methods^6–10^ take a direct approach at these questions: The idea of state-based methods is that each macromolecular configuration is mapped to a single global state, typically such that the state distinguishes between metastable states – sets of configurations that are separated by rare-event transitions. The number of states of a macromolecule is then determined by the timescales one wants to resolve^2^, and an MD simulation can be considered converged when all these metastable states have been found and the transitions between them have been sampled in both directions. This state-based view has led to the extensive characterization of folding^11–13^, conformation changes^14,15^, ligand binding^16–19^ and association/dissociation^20^ in small to medium-sized proteins. Although seemingly conceptually different, reaction-coordinate (RC) based methods, such as umbrella sampling^21^, flooding/metadynamics^22,23^ and related analyses are also state-based methods, where the state is characterized by the values of the chosen RCs. Still, with such methods, all rare events that are not statistically independent of the chosen RCs must be sampled, and the corresponding metastable states resolved.

Even with massive simulation power and enhanced sampling methods, a converged analysis in the state-based picture fundamentally relies on the fact that there are relatively few metastable states. This is the case for cooperative macromolecules, such as small to medium-sized proteins, where the long-ranged interactions create relatively smooth free energy landscapes, explaining the success for these systems. However, almost all available MD simulation and analysis methods will break down for systems with many metastable states. The proliferation of metastable states can already be observed in nontrivial protein systems^24^, but a more pathological example are nucleic acids where the possible ways to pair bases whose number grows exponentially with system size creates highly rugged energy landscapes with many metastable states^25,26^. Even for protein systems, the ability to characterize the global system state by a single variable will break down at a certain size. Beyond the size of a few nanometers, electrostatic interactions are weak, and thus large protein machines such as the ribosome, the spliceosome and neuronal active zones are expected to consist of dozens to thousands of largely independently switchable units. Even if each of these units only has two possible states, the total number of global system states grows exponential with system size, and any simulation or analysis method that relies on all these states to be sampled is doomed.

In this paper, we propose a change of perspective by introducing Dynamic Graphical Models (DGMs) that characterize each molecular configuration by a vector of substate configurations. DGMs then model how these substates evolve in time by a set of local rules, each of which govern the dynamics of each substates in the field of the others. Although conceptually less simple than models with a single global state, such as MSMs, the decision to model a system by many coupled substates is key in reducing the computational complexity for systems that have astronomically large numbers of states. In order to learn the rules according to which each subsystem *i* switches its state, not all global system configurations need to be sampled, but only the the states of those other subsystems *j* that are directly coupled to *i*. This idea is conceptually similar to a recent Granger causality model which captures the time-evolution of several binary random variables^27^. DGMs however build upon the idea of Ising models which are graphical models used to model key phenomena in wide array of disciplines^28–30^. The main difficulty in applying these ideas to macromolecular dynamics is that it is not *a priori* clear what the subsystems are and how they are coupled. Estimating the structure of graphical models from data is an extensively studied machine learning problem^31–34^. Here we take a first step towards estimating DGMs from simulation data by employing a dynamical version of Ising model estimators^35^.

The property that not all global system configurations need to be sampled to parametrize a DGM implies that DGMs are generative models. That means, that they should be able to predict previously unobserved molecular configurations and thus promote the efficient discovery of conformation space. Here we demonstrate that DGMs can indeed predict previously unobserved protein conformations and that protein thermodynamics and kinetics can be modeled in a similar way as with MSMs.

## Dynamic Graphical Models

### General Model Structure

Here, we propose Dynamic Graphical Models (DGM) as a new method to model molecular kinetics and thermodynamics, inspired by Graphical Models, a classical Machine Learning approach to encode dependencies between random variables^33^. Instead of encoding the global molecular configuration in a single state variable, DGMs characterize the molecular configuration by a set of *sub-systems* 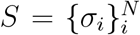 that encode the states of local molecular features (Fig. 1). The *coupling parameters J_ij_*(*τ*) of a DGM then encode how each subsystem switches in time as a result of the states of other subsystems at a previous time, i.e. *σ_i_*(*t*) *→ σ_j_*(*t* + *τ*). Like MSMs, DGMs are Markovian models in which the current configuration only depends on the previous time step. In contrast to MSMs that encode transition probabilities of global system configurations, DGMs encode the transition probabilities of single subsystems, dependent on the settings of all the subsystems. In addition, each subsystem does generally not depend on the all other subsystems, but on a local neighborhood of subsystems – in other words the coupling parameters are generally sparse.

**Figure 1.**
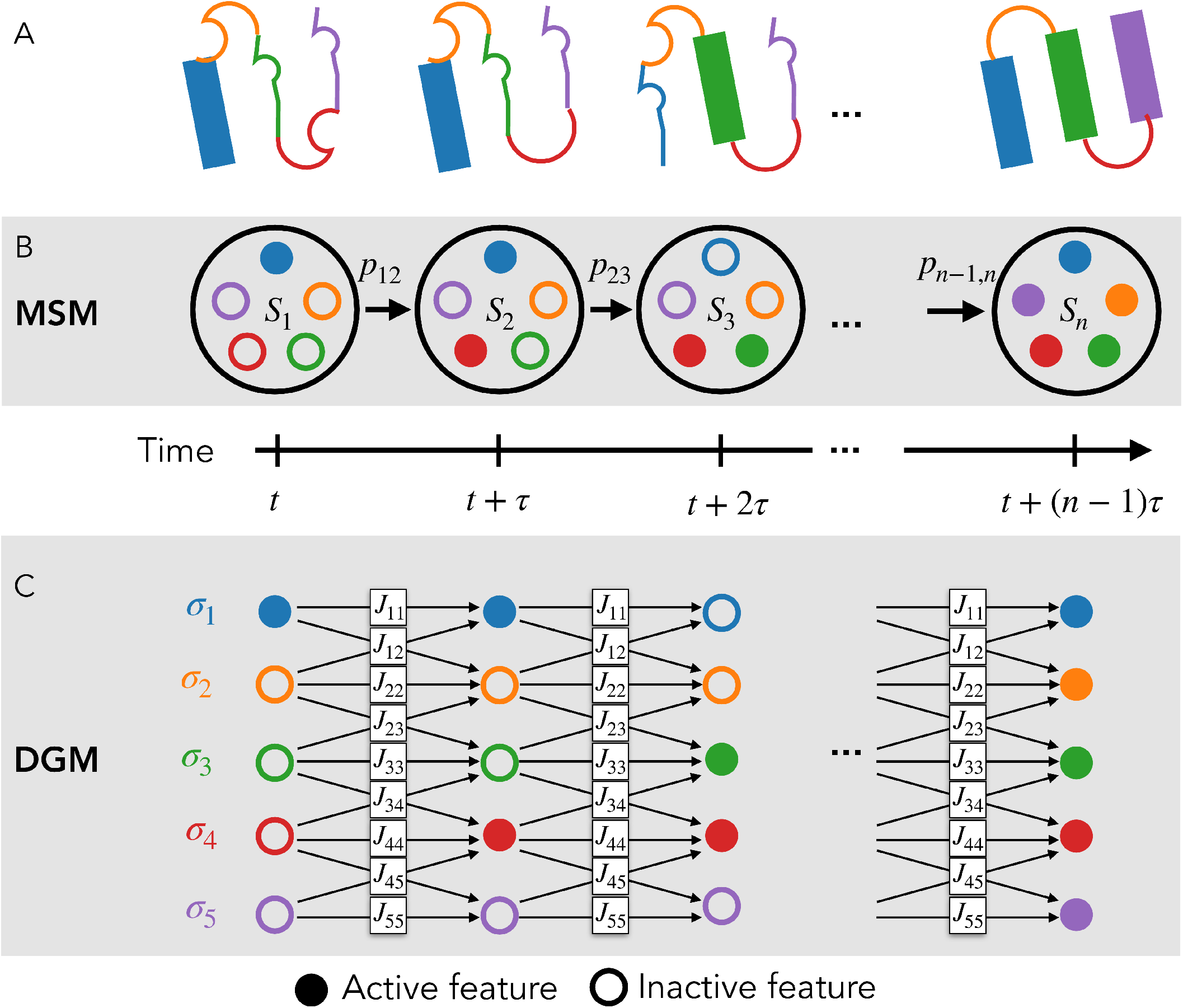
Illustration of molecular representations in Markov state models (MSM) and dynamic graphical models (DGM). A: cartoon representation of protein conformational states found along a trajectory. B: MSM with Markov states, *S_i_*, and transition probabilities between states *S_i_* and *S_j_*, *p_ij_*. The number of diσerent Markov states, *S_i_*, may grow exponentially with the number of features, and only feature combinations that have been observed can be encoded in Markov states. C: DGMs represents the current state of the system via the states of its sub-systems, *σ_i_* that are coupled by parameters *J_ij_*. The DGM can still encode exponentially many states, but the number of model parameters grows much slower. DGMs can predict system states that have not yet been observed.

For illustration, consider a DGM of a one-dimensional Ising model^36^ (Fig. 2A) consisting of *N* sub-systems on a ring, each of which adopt one of two configurations, 1 or *−*1. Each sub-system (or spin) interacts directly only with its nearest neighbors, and possibly an external field, resulting in three coupling parameters for each spin. Given a choice of the dynamics used to flip the spins, such as Metropolis, Glauber (Gibbs) or Kawasaki dynamics^37–39^, a DGM that describes the probability of each spin’s state at time *t* + *τ* given the spin configurations at time *t* will thus require 3*N* parameters. A direct MSM, on the other hand, distinguishes all 2^*N*^ global Markov states and needs to estimate an up to 2^*N*^ × 2^*N*^ transition matrix from the data. More importantly, an MSM can only retrospectively describe the dynamics between states that have been observed in the simulation data. For example, an MSM is unable to predict that “spins up” and “spins down” are two different metastable states unless it has sampled transitions between them. A DGM can in principle be parameterized only using the spin fluctuations in one of the two metastable states, and the global thermodynamics and kinetics can still be predicted. We intend to exploit this property for modeling biomolecular dynamics.

**Figure 2.**
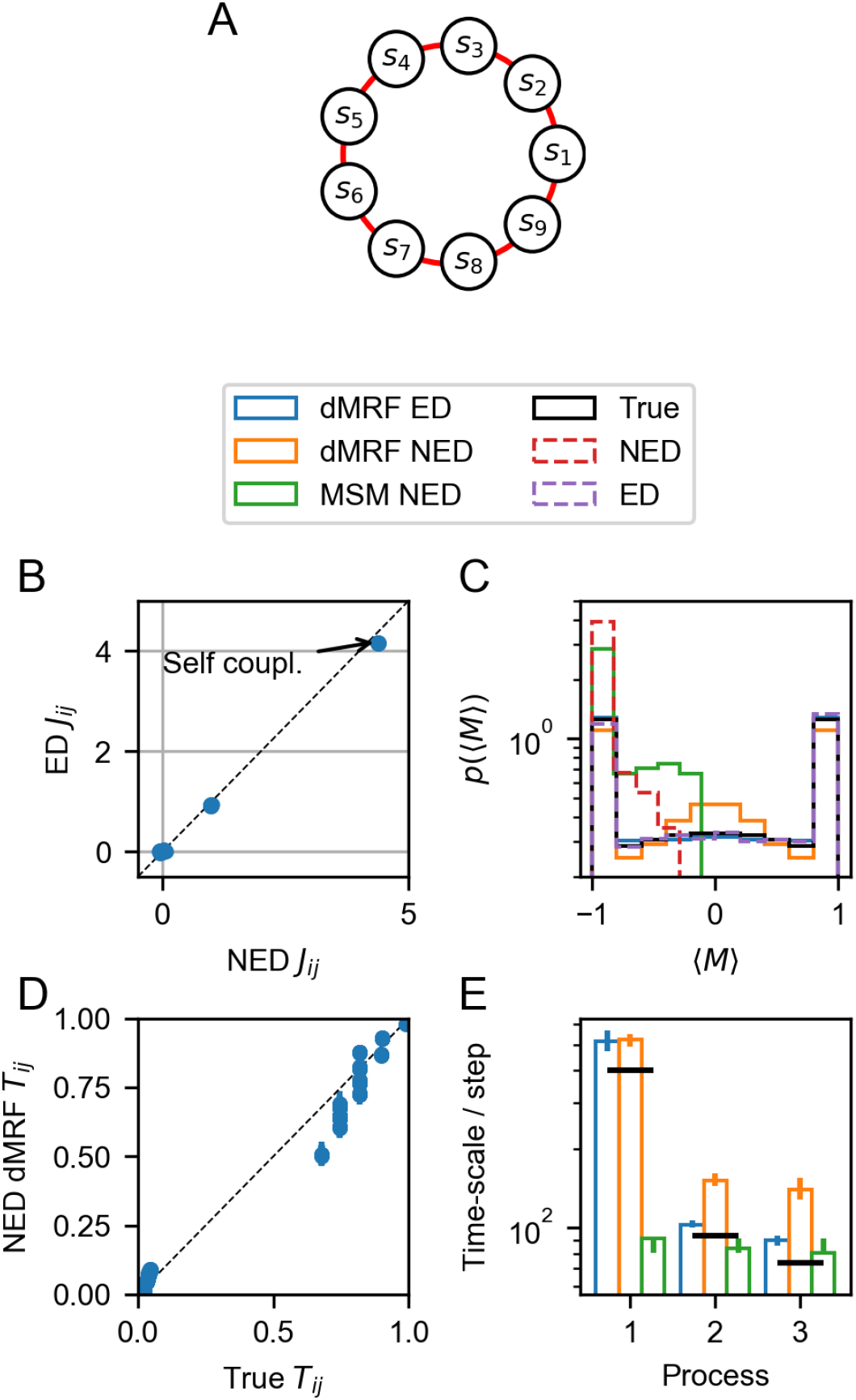
DGMs estimated for a one-dimensional Ising model from equilibrium simulation data (ED) and from simulation data only containing configurations with negative net magnetization (NED). A) Illustration of the 9-spin 1D Ising model with periodic boundary conditions. B) Comparison of coupling parameters estimated using equilibrium and non-equilibrium data sets. C) Analytic stationary distributions of〈*M*〉 and empirical histograms of ED and NED data sets. Shown DGM predictions are for models where the local fields {*h_i_*} are not estimated. D) Comparison of global configurationsal state transition probabilities predicted in DGM trained on non-equilibrium data set against the true reference values.E) Implied time-scales of estimated models and the true reference.

When using DGMs to model biomolecular dynamics, we are not given the definition of subsystems and their couplings a priori, but we must rather estimate them from data. A sub-system could be something as complex as a protein domain (Fig. 1A) with multiple internal states, or something as simple as a torsion angle rotamer or a contact between two chemical groups that each have only two settings, similar as spins. Once the sub-systems are defined, it must be estimated which of them are coupled. Estimating the coupling graph from data is a notoriously difficult problem for graphical models and the related Markov Random Fields (MRF)^31,32,40^. However, we have the advantage that modeling the dynamics via a DGM creates a directed dependency graph in time: the spin variables at time *t* + *τ* only depend on the spin variables at time *t* (Fig. 1C). This makes the problem of estimating the coupling between sub-systems tractable.

### Dynamic Ising Models

Here we propose a first implementation of DGMs that is based on the idea of Ising models. Ising models consist of a set of *N* spins with configurations 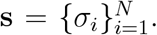 Each spin can be in one of two settings, *σ_i_* ∈ {−1, 1}. The equilibrium probability of a spin configurations is given by:

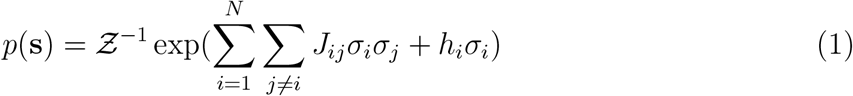

where the (unit-less) model parameters *J_ij_* = *J_ji_* and *h_i_* describe the coupling strength between sub-systems *i* and *j* and the local field of sub-system *i*, respectively, and *Ƶ* is the partition function. The most common Ising models have spins arranged in a lattice and only neighboring spins are coupled, however the general Ising model (1) allows arbitrary spin couplings and does not prescribe a specific topology. For given parameters {*J_ij_*} and {*h_i_*}, the distribution (1) can be easily sampled^37,38^.

When modeling molecular kinetics our data consists of (possible short) time-series rather than samples from the equilibrium distribution. Therefore, we employ Dynamic Ising Models (DIMs)^35^ that model the conditional distribution *p*(s_*t*_ ~ s_*t−τ*_) that governs how spin configurations – and thus the configurations of our molecular system – changes in time. Note that given a model *p*(s_*t*_ ~ s_*t−τ*_) we can simulate long trajectories and thus also predict the equilibrium distribution, *p*(s). The DIM model is given by:

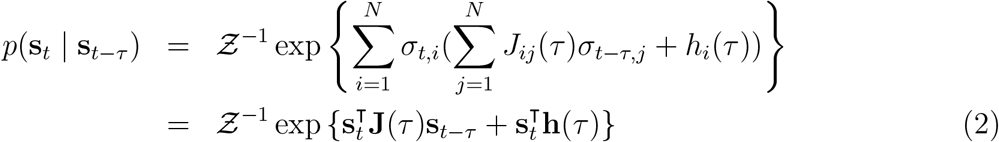

where **J**(*τ*) ∈ ℝ ^*N*×N^ is a matrix of coupling parameters and **h**(*τ*) is a vector of local fields and *Ƶ* is the partition function. DIMs can readily be generalized more than two states per subsystem *σ_i_* (see SI for details)

### Properties and Estimation of Dynamic Ising Models

Unlike in eq. (1), the self-couplings *J_ii_*(*τ*) may be nonzero and the coupling matrix **J**(*τ*) is not necessarily symmetric (*J_ij_*(*τ*) = *J_ji_*(*τ*)).

In Glauber dynamics only a single sub-system is allowed to change its state in some infinitesimal time *δt*, in contrast, the DIM model samples the current state of all sub-systems given their configurations at a time *τ* prior. Consequently, if we use a *τ* which is larger than the characteristic time-scale of one or more of the sub-systems, the dynamics of these subsystems will not be resolved. This is analogous to how a lag-time defines the temporal resolution in a Markov State Model. The relationship between DIM model parameters and the macroscopic system properties, such as relaxation rates, are further analyzed in the Supporting Information and Fig. S1.

The problem of estimating eq. (2) from time-series data has previously been investigated in the context non-equilibrium network reconstruction^35^. We can exploit that the problem of estimating Ising model parameters can be cast as a set of *N* logistic regression problems. Here, we generalize this method by showing that a DIM with more than two states per subsystem (which may be called a Dynamic Markov Random Field or Dynamic Potts Model) can solved by softmax regression (SI for details).

In order to illustrate DIM estimation, consider the conditional probability of a single sub-system *σ_t,i_*, given s_*t−τ*_

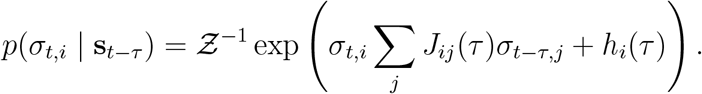

Recall, that *σ_t,i_* can only take on two configurations in the Ising model {*−*1, 1}. This means that we can evaluate the partition function *Z* analytically as it is the sum of the only two outcome probabilities, *p*(*σ_t,i_* = 1 *~* s_*t−τ*_) and *p*(*σ_t,i_* = *−*1 *~* s_*t−τ*_). That is, the partition function for the single sub-system conditional transition probability is independent of the number of global system configurations, in contrast to the equilibrium formulation of the Ising model (eq. (1)) where computation of *Ƶ* requires the summation of all global configurations of the system. These observations make the DIMs computationally attractive as exact calculation of *Ƶ* is tractable even for systems with a large number of sub-systems. Using this we may realize that *p*(*σ_t,i_ ~* s_*t−τ*_) follows a sigmoid in *θ_t–τ.i_* = Σ*_j_ j_ij_*(*τ*)*σ_t–τ,j_* + *h_i_*

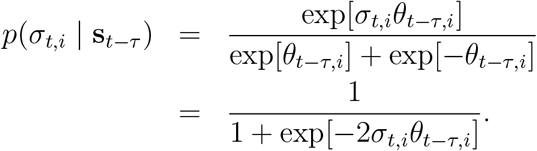

Consequently, estimating and 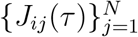 and *h_i_* corresponds to estimating the *i*’th row in the coupling matrix **J**, and the *i*’th element of the field vector **h**.

Since the marginal probability distributions of the sub-systems of s_*t*_ are conditionally independent given s_*t−τ*_, the joint probability density is simply their product,

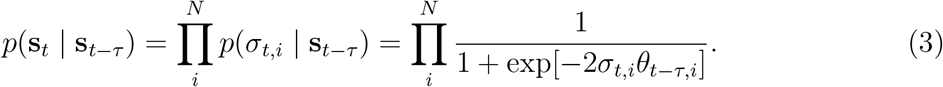

which facilitates the DIM estimation to be solved by performing an independent logistic regression problem for each sub-system.

Overall, given a time-series of state-configurations, S = {s_0_, s_*τ*_, …, 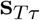}, the DIM parameters can be found by maximizing the likelihood function:

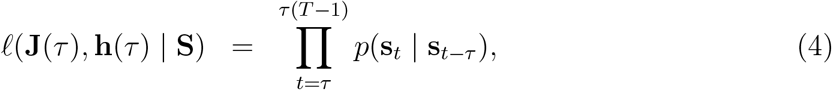

If we combine this likelihood with a Laplacian prior, maximizing the posterior

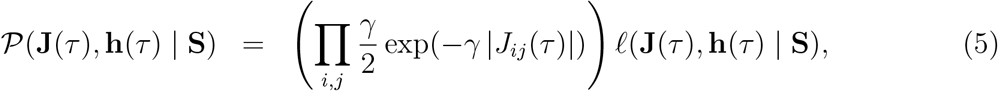

favors sparse solutions with some *J_ij_* = 0, where the regularization parameter γ controls the degree of sparsity.

Here, Eq. (5) is optimized using the SAGA optimization algorithm^41^, as implemented in the scikit-learn python library^42^.

As part of this manuscript we have developed a library for estimation and analysis of DGMs which along with example notebooks is available at: http://www.github.com/markovmodel/graphtime/

## Results

### Recovery of an unobserved phase, its thermodynamics and kinetics in the Ising model from biased data

One of the most interesting properties for DGMs is that they make predictions for global system configurations that have not been observed in the training data. Here we test whether an Ising model DGM trained only from data with negative net magnetization can predict the existence of configurations with positive net magnetization, their equilibrium probabilities and the timescale of remagnetizing the system.

As a first test, we simulate a one-dimensional periodic Ising model with 9 spins using Glauber dynamics which exhibits two metastable phases with negative and positive net magnetization, 〈*M*〉. Non-equilibrium (NED) and equilibrium (ED) data sets are generated by simulating 16 trajectories with all sub-systems initialized in state *−*1. NED simulations are terminated before the net magnetization becomes larger than 0, i.e., the NED data set has no configurations with more than 4 sub-systems in the +1 configurations. Consequently, the entire metastable state with positive net magnetizations is missing. In the ED data simulations are run with a length so as to match the sampling statistics of the NED data, but trajectories are allowed to reach positive net magnetizations (Fig 2A). DGMs are estimated by optimizing Eq. (5). We find that both the ED and NED DGMs converge to the correct sub-system couplings (Fig 2B) regardless of whether we choose to estimate the external field parameters, *h_i_*, or not (Fig S2). This means that the unbiased sub-system couplings can be recovered from a biased data-set, in turn suggesting that global system characteristics can be estimated from incomplete data.

In order to compute the overall thermodynamics and kinetics of the system and the estimated DGMs, we generate MSMs with all 2^9^ states and compute their transition probabilities using eq. (3). Indeed, the distribution of the net magnetizations, 〈*M*〉, predicted by the DGMs closely resemble the true distribution (Fig 2C). In contrast, a MSM estimated using the NED data only contains configurations contained in the data and therefore fails to predict the existence of positive net magnetizations (Fig 2C).

Remarkably, DGMs can also predict the kinetics involving states that have not been observed. The DGM predictions for the 2^9^*×*2^9^ transition probabilities between global system configurations correlate well with their true values, albeit with some biases (Fig 2D). The true global relaxation time-scales are recapitulated well by the DGM models (Fig 2E) – however, for the NED DGM the time-scales are systematically overestimated, as some self-couplings are over-estimated (Fig 2B). As expected, the NED MSM completely fails to capture the slowest relaxation timescale as it corresponds to the inversion of net magnetization which has not been observed in the training data.

### DGMs as models of molecular dynamics

We now turn away from the “ideal” case of a discrete, Markovian system to Molecular Dynamics (MD), where DGMs are build upon a discretization of molecular features and hence introduce a systematic modeling error. To this end, we model MD simulation data of a small penta-peptide system (WLALL) and compare DGMs with MSMs that are well established for peptide simulation data (Fig. 3A).

**Figure 3.**
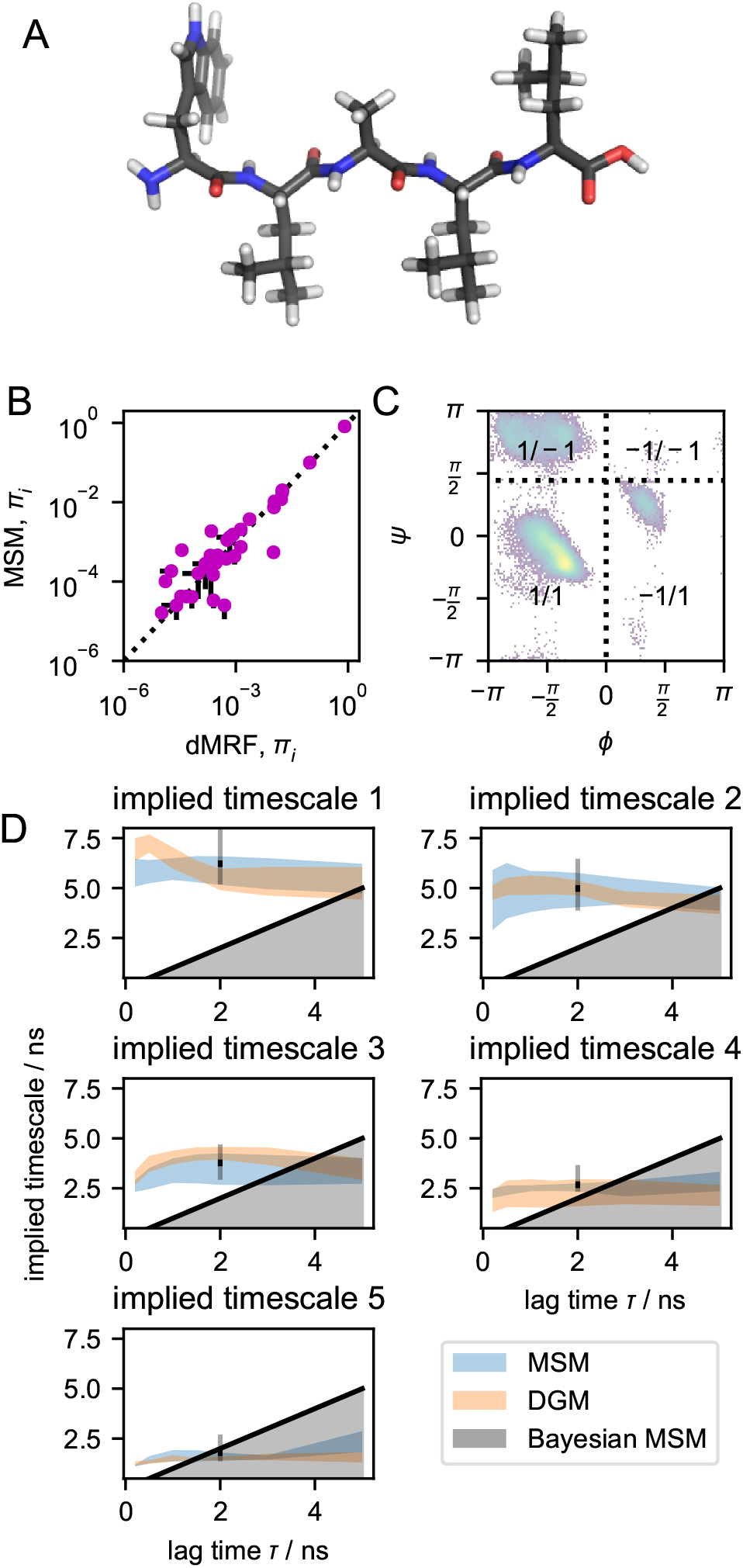
Comparison of DGM and MSM describing back-bone torsion dynamics of the pentapeptide Trp-Leu-Ala-Leu-Leu (WLALL). A) Molecular structure render of the WLALL peptide in stick representation. B) Global state distribution computed according to DGM and MSMs. C) Torsion angles serve as binary subsystems whose value indicate the rotamer. Vertical and horizontal dotted lines indicate the boundaries of the −1 to 1 transition in the *ϕ* and *ψ* torsions, respectively. Translucent reference density is a histogram of (*ϕ,ψ*,*ψ*) torsion pairs in Leu-2 from the WLALL peptide simulation dataset. D) 95% confidence intervals of MSM (orange) and DGM (blue) implied time-scales. Confidence intervals in B and D are computed using bootstrap with replacement. Markov states shown in B are those observed in all 24 trajectories; DGM state probabilities are renormalized to this set.

We model WLALL by seven sub-systems corresponding to its back-bone torsions (*ϕ,ψ*). Each subsystem has two states, defined by splitting *ϕ* at 0° and *ψ* at 80° (SI), thus discretizing the Ramachandran plane into four states (Fig. 3C). We estimate MSMs and DGMs using this representation. For the MSMs, all 2^7^ possible discrete angle configurations serve as possible Markov states and the MSMs are estimated on the subset of observed Markov states. DGMs are estimated by maximizing eq. (5), and, for comparison, translated into MSMs by enumerating all possible global state transitions. We find that the implied time-scales computed from DGMs match those of the MSMs well at multiple lag-times, within statistical error (Fig. 3B). Similarly, the stationary probabilities of the Markov states match closely, in particular for high probability states (Fig. 3C).

The DGMs only use 7^2^ + 7 = 72 parameters but predict equilibrium probabilities for all 2^7^ = 128 possible states and all 2^14^ = 16, 384 possible transitions. Thus, DGMs statistically much more efficient than MSMs. MSMs directly estimate a transition matrix for all states that have been observed – for WLALL 45 states (35.16%) and 337 state-to-state transitions (2.06%) have been observed (*τ* = 2 ns). Enumerating all possible system configurations is not scalable to larger systems. For larger systems, MSMs are built by clustering states, preferentially by grouping states that are quickly exchanging^43–46^. Still, this approach requires all long-lived sets of configurations to be sampled in the data. Subsequently, we show that DGMs can predict unobserved metastable states for larger biomolecules and therefore systematically outperform MSM approaches to data analysis.

### Prediction of unobserved meta-stable molecular configurations

To more systematically test the predictive power of DGMs in predicting meta-stable states not observed in the simulation data we estimated several DGMs designed to be selectively blind towards a particular meta-stable state. We used data of two fast-folding proteins previously published, the *α*-helical protein villin and the *α* / *β* protein BBA^47^.

For villin and BBA we built a hidden Markov model (HMM)^48^ that resolves five and a four metastable states, respectively (structures in Fig. 4A/E). We then estimated a DGM using the same data and confirmed that it reproduces the equilibrium probabilities of the HMM metastable states (Fig. 4B/F, last panels). Using the HMM assignments of MD data to meta-stable states, we generate five and four sets of training data for the two proteins, where each set is missing one of the metastable states. These sub-sampled data-sets are used for estimating “one-blind” DGMs that are each “blind” to one meta-stable configurations (see SI for details). From the estimated DGMs we simulate synthetic trajectories of the time-evolution of the sub-systems, in order to test whether these one-blind DGMs can recover the unobserved states and their statistics.

**Figure 4.**
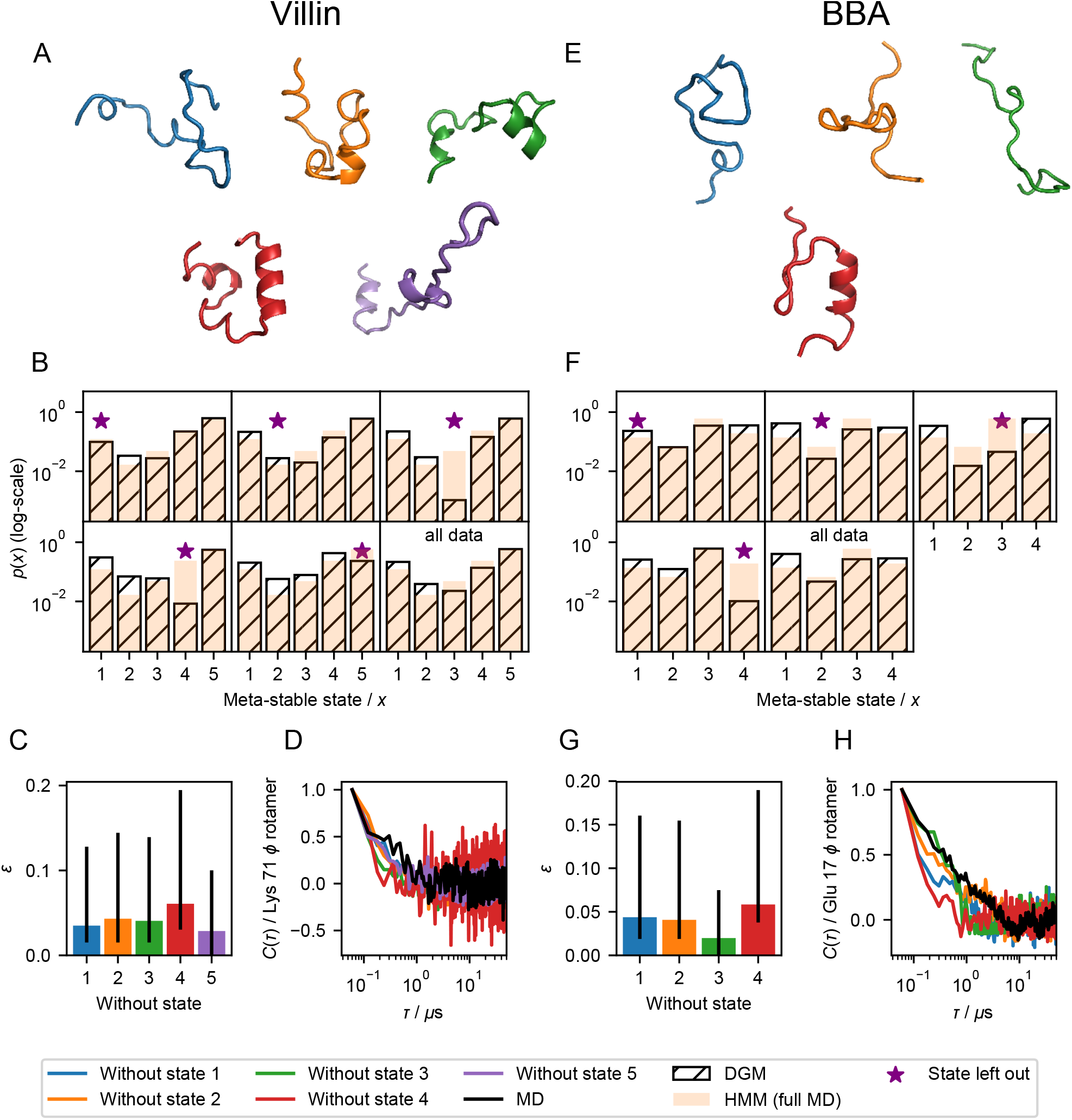
Prediction of macroscopic stationary and dynamic properties of fast folding proteins Villin and BBA with DGMs. Molecular renders of meta-stable configurations identified by HMMs of villin (A) and BBA (E), using a ribbon representation. Color/meta-stable state relationship: Blue/1, Orange/2, Green/3, Red/4 Purple/5. B/F) Meta-stable state distributions sampled by DGMs estimated leaving data from one of four meta-stable state out during estimation and using the full data set (hatched). Reference distribution estimated using a hidden Markov model (HMM) estimated with the full MD data-set (orange). C/G) Bar plots of fraction incorrectly assigned sub-systems (*ψ*) of atomistic models from sub-system encoded trajectories sampled by DGMs, errorbars represent a 68% confidence interval. D) Normalized auto-correlation function of the rotameric state of the *ϕ* torsion of Lys 71 in Villin as sampled by the DGM models and in the simulation data. H) Normalized auto-correlation function of the rotameric state of the *ϕ* torsion of Glu 17 in BBA as sampled by the DGM models and in the simulation data.

We find that the one-blind DGMs generally sample the same configurationsal space as the full MD simulations, although the predicted equilibrium distributions somewhat differ (Fig. S3). This suggests that although the predicted thermodynamics in the one-blind DGMs is not quantitatively accurate, they are able to predict relevant states that have not been included in the training data. At a more microscopic level this is reflected by the fact that statistical properties of the configurations sampled by the DGM closely agree with their MD counterparts (Figs. S4, S5 and 4C/G).

Next we compare the macroscopic stationary and dynamics properties predicted by the one-blind DGMs with the full-data HMMs (Fig. 4). This comparison confirms that all one-blind DGMs are able to predict the existence of the state that has been missing in the training data, and in most cases with a population comparable to the MD simulation (Fig. 4B/F). For visualization we have reconstructed molecular trajectories from the sub-system trajectories simulated from a select set of one-blind DGMs (Suppl. Vids. 1,2).

Finally, we compare the kinetics predicted by the one-blind DGMs with those of the original simulation. As an example, we compute the time-correlation function (TCF) of the rotameric state of a backbone torsion angle in the reference simulations of villin and BBA and compare them to corresponding TCFs from the DGM trajectories (Fig. 4D/H). We find these generally agree well, although the DGMs tend to somewhat over-estimate the relaxation rates, in particular the DGMs that were blind to the folded configurations (4th (red) state in both cases).

### Meta-stability in one-blind DGMs

The analysis above shows that the conformational space sampled by the one-blind DGMs is realistic and recapitulates the conformational space sampled in the full MD datasets. However, whether meta-stable states predicted by the one-blind DGMs indeed recapitulate the metastable states identified in the MD is not clear from this analysis alone. To more quantitatively assess this, we estimated HMMs trained on data-sets generated using the one-blind DGMs (SI). In half of the cases the number of meta-stable states identified from the one-blind DGM datasets matches the corresponding numbers of from the full MD dataset (Table S1). In most of the remaining cases one meta-stable state is missing, and in a single case one extra meta-stable state is identified, closely resembling another meta-stable state (SI).

To better understand the identity of the meta-stable states observed in the one-blind DGMs we assigned them to corresponding meta-stable states from the MD dataset such as to minimize the total assignment error (SI). We find that the meta-stable states sampled by the one-blind DGMs can be unambiguously assigned to a corresponding meta-stable states observed in the MD dataset, and their sampling statistics correlate well with their MD counterparts (Fig. S7A,C). The average lifetimes of these meta-stable states qualitatively correlate with those of the reference MD dataset, but generally are too short, as already indicated above. (Fig. S7B,D).

## Discussion and Conclusions

The ability to simulate a molecular system to convergence depends critically on how the simulation is analyzed and how the quantities of interest are computed. Most current simulation and analysis methods employ the idea that the molecule is a single global state – most explicitly, MSMs make that assumption. However, this picture leads to a sampling problem that does not scale to large molecular systems. A molecule with *N* independent subunits, each with *M* metastable states each will have an exponential number, *M^N^*, of metastable states.

In this paper, we introduce the concept of dynamic graphical models (DGM) and their use to model molecular kinetics. In DGMs, the global configurationsal state is encoded as a vector of states of its sub-systems. As DGMs only model the pair-wise interactions between subsystems an exponential number of global configurations can be represented with a quadratic number of parameters. We illustrate dynamic graphical models using the Dynamic Ising Model — and its generalization, the Dynamic Markov Random Field (DMRF, in SI).

In general, we find the DGMs, in spite of their compact parametric form can approximate well the dynamic behavior of several molecular systems. When gauged against MSMs, DGMs predict similar characteristic time-scales and stationary distributions. Beyond the capabilities of MSMs, DGM also allow for the prediction of transitions to and from unobserved molecular conformations and their stationary probabilities. Specifically, we found that DGMs can predict the predict the presence of meta-stable configurations which have not been observed during model estimation, although their absolute probabilities are not always accurately captured.

The new representation of a molecule in the DGM framework gives rise to new technical challenges that need to be addressed in future studies. (1) How can the sub-systems of a molecule be learned from simulation data? Here we have defined these sub-systems *a priori*, e.g., by choosing dihedral rotamers, but general optimization principles are needed, e.g. by exploiting the variational approach for Markov processes (VAMP)^49–51^. (2) Going from an encoded representation back to a molecular configurations (decoding) is generally a difficult problem. (3) Although efficient regularized maximum likelihood estimators are readily available, optimally choosing regularization factors remains heuristic. (4) The compact parametric form of the DGMs comes with limited expressive power. It is specifically assumed that the sub-system couplings are *not* a function of the global configurations – if this is violated we cannot expect to obtain quantitatively predictive models using DGMs. (5) Unlike for MSMs^52^, the integration of experimental data into the estimation of DGMs is currently not possible. While in MSMs, the experimental observable can be directly expressed in terms of transition matrix properties^53,54^, in DGMs it involves sampling which could be both time-consuming and difficult, in particular when no sub-system decoding is possible.

Machine learning, and in particular deep learning, has seen a surge in interest in the sciences the past few years, including in the context of molecular kinetics^55,56^. A hallmark of deep learning is its ability to learn complicated non-linear functions which fulfill a prespecified set of criteria given suficient available data^57^. The identification of sub-systems and their discretization is highly non-linear and akin to the featurization, projection and discretization process whose success critically determines the success of Markov state modeling. The VAMPnets approach recently illustrated how the featurization, projection and discretization pipeline of MSM building could be replaced by a deep neutral network^55^. We envision that a related approach can be taken in the context of identifying sub-systems in molecular systems for DGMs and thereby possibly side-step this currently cumbersome and manual process.

We anticipate DGMs and follow-up methods to be broadly applicable in biophysics. The compact parametric form allows us to model molecular systems which are significantly larger than what would be tractable using for example MSMs. Furthermore, a sparse sub-system coupling matrix **J**(*τ*) could enable a spatial decomposition or large molecular systems into conditionally independent fragments, more amenable for simulation. The ability of DGMs to predict unobserved global configurations could further be used in an adaptive simulation setting, where previously unobserved states are reconstructed and used to seed new molecular simulations.

**Synopsis** Discovery of meta-stable configurationsal states of large biomolecules is an important problem in biophysics. We describe a procedure to speed up this discovery process by learning how local structural elements interact and how these interaction influence the temporal behavior of the global configurationsal states.

### Supporting Information Available

A detailed report of the practical estimation of models presented in the work including hyper-parameter choices can be found in the supporting information. Further, a theoretical generalization of DGM to cases where sub-system may adopt more than two discrete states along with their constrained and unconstrained maximum likelihood estimators is available. Finally auxiliary results including figures and movies illustrating properties of the estimated models may are shown in the supporting material. This material is available free of charge via the Internet at http://pubs.acs.org/.

## Acknowledgement

We thank Fabian Paul, Hao Wu, Christoph Wehmeyer, Illia Horenko, Moritz Hoffmann and Tim Hempel for stimulating discussions. We thank Nuria Plattner for providing simulation data for the penta-peptide system and DE Shaw Research for providing the simulation data for the villin headpiece and BBA. We gratefully acknowledge funding from the Alexander von Humboldt foundation (Postdoctoral fellowship to SO) and the European Research Council (ERC Consolidator Grant ScaleCell to FN).

## References

(1) Schütte, C.; Fischer, A.; Huisinga, W.; Deuflhard, P. J. Comput. Phys. 1999, 151, 146–168.

(2) Noé, F.; Horenko, I.; Schütte, C.; Smith, J. C. J. Chem. Phys. 2007, 126, 155102.

(3) Chodera, J. D.; Dill, K. A.; Singhal, N.; Pande, V. S.; Swope, W. C.; Pitera, J. W. J. Chem. Phys. 2007, 126, 155101.

(4) Buchete, N. V.; Hummer, G. J. Phys. Chem. B 2008, 112, 6057–6069.

(5) Prinz, J.-H.; Wu, H.; Sarich, M.; Keller, B. G.; Senne, M.; Held, M.; Chodera, J. D.; Schütte, C.; Noé, F. J. Chem. Phys. 2011, 134, 174105.

(6) Du, W.-N.; Marino, K. A.; Bolhuis, P. G. J. Chem. Phys. 2011, 135, 145102.

(7) Noé, F.; Krachtus, D.; Smith, J. C.; Fischer, S. J. Chem. Theo. Comp. 2006, 2, 840–857.

(8) Preto, J.; Clementi, C. Phys. Chem. Chem. Phys. 2014, 16, 19181–19191.

(9) Singhal, N.; Pande, V. S. J. Chem. Phys. 2005, 123, 204909.

(10) Rohrdanz, M. A.; Zheng, W.; Maggioni, M.; Clementi, C. J. Chem. Phys. 2011, 134, 124116.

(11) Noé, F.; Schütte, C.; Vanden-Eijnden, E.; Reich, L.; Weikl, T. R. Proc. Natl. Acad. Sci. USA 2009, 106, 19011–19016.

(12) Bowman, G. R.; Voelz, V. A.; Pande, V. S. J. Am. Chem. Soc. 2011, 133, 664–667.

(13) Zheng, W.; Qi, B.; Rohrdanz, M. A.; Caflisch, A.; Dinner, A. R.; Clementi, C. J. Phys. Chem. B 2011, 115, 13065–13074.

(14) Kohlhoσ, K. J.; Shukla, D.; Lawrenz, M.; Bowman, G. R.; Konerding, D. E.; Belov, D.; Altman, R. B.; Pande, V. S. Nat. Chem. 2014, 6, 15–21.

(15) Bowman, G. R.; Bolin, E. R.; Harta, K. M.; Maguire, B.; Marqusee, S. Proc. Natl. Acad. Sci. USA 2015, 112, 2734–2739.

(16) Buch, I.; Giorgino, T.; De Fabritiis, G. Proc. Natl. Acad. Sci. USA 2011, 108, 10184–10189.

(17) Plattner, N.; Noé, F. Nat. Commun. 2015, 6, 7653.

(18) Gu, S.; Silva, D.-A.; Meng, L.; Yue, A.; Huang, X. PLoS Comput. Biol. 2014, 10, e1003767.

(19) Huang, D.; Caflisch, A. PLoS Comput Biol 2011, 7, e1002002+.

(20) Plattner, N.; Doerr, S.; Fabritiis, G. D.; Noé, F. Nat. Chem. 2017, 9, 1005–1011.

(21) Torrie, G. M.; Valleau, J. P. J. Comp. Phys. 1977, 23, 187–199.

(22) Grubmüller, H. Phys. Rev. E 1995, 52, 2893.

(23) Laio, A.; Parrinello, M. Proc. Natl. Acad. Sci. USA 2002, 99, 12562–12566.

(24) Paul, F.; Wehmeyer, C.; Abualrous, E. T.; Wu, H.; Crabtree, M. D.; Schöneberg, J.; Clarke, J.; Freund, C.; Weikl, T. R.; Noé, F. Nat. Commun. 2017, 8, 1095.

(25) Thirumalai, D.; Woodson, S. A. Acc. Chem. Res. 1996, 29, 433–439.

(26) Keller, B. G.; Kobitski, A. Y.; Jäschke, A.; Nienhaus, U. G.; Noé, F. J. Am. Chem. Soc. 2014, 136, 4534–4543.

(27) Gerber, S.; Horenko, I. Proceedings of the National Academy of Sciences 2014, 111, 14651–14656.

(28) Yap, W.; Saroσ, H. Journal of Theoretical Biology 1971, 30, 35–39.

(29) Schneidman, E.; Berry, M. J.; Segev, R.; Bialek, W. Nature 2006, 440, 1007–1012.

(30) Stauσer, D. American Journal of Physics 2008, 76, 470–473.

(31) Ravikumar, P.; Wainwright, M. J.; Laσerty, J. D. Ann. Statist. 2010, 38, 1287–1319.

(32) Parise, S.; Welling, M. In Advances in Neural Information Processing Systems 19; Schölkopf, B., Platt, J., Hoσman., T., Eds.; 2006.

(33) Bishop, C. M. Pattern Recognition and Machine Learning; Springer Science Business Media, 2006.

(34) Koller, D.; Friedman, N. Probabilistic Graphical Models: Principles and Techniques (Adaptive Computation and Machine Learning); The MIT Press, 2009.

(35) Roudi, Y.; Hertz, J. Physical Review Letters 2011, 106.

(36) Lenz, W. Phys. Z. 1920, 21, 613–615.

(37) Glauber, R. J. Journal of Mathematical Physics 1963, 4, 294–307.

(38) Kawasaki, K. Physical Review 1966, 145, 224–230.

(39) Metropolis, N.; Rosenbluth, A. W.; Rosenbluth, M. N.; Teller, A. H.; Teller, E. The Journal of Chemical Physics 1953, 21, 1087–1092.

(40) Roudi, Y.; Tyrcha, J.; Hertz, J. Physical Review E 2009, 79.

(41) Defazio, A.; Bach, F.; Lacoste-Julien, S. In Advances in Neural Information Processing Systems 27; Ghahramani, Z., Welling, M., Cortes, C., Lawrence, N. D., Weinberger, K. Q., Eds.; Curran Associates, Inc., 2014; pp 1646–1654.

(42) Pedregosa, F. et al. Journal of Machine Learning Research 2011, 12, 2825–2830.

(43) Molgedey, L.; Schuster, H. G. Physical Review Letters 1994, 72, 3634–3637.

(44) Ziehe, A.; Müller, K.-R. ICANN 98; Springer London, 1998; pp 675–680.

(45) Perez-Hernandez, G.; Paul, F.; Giorgino, T.; D’Fabritiis, G.; Noé, F. J. Chem. Phys. 2013, 139, 015102.

(46) Schwantes, C. R.; Pande, V. S. Journal of Chemical Theory and Computation 2013, 9, 2000–2009.

(47) Lindorσ-Larsen, K.; Piana, S.; Dror, R. O.; Shaw, D. E. Science 2011, 334, 517–520.

(48) Noé, F.; Wu, H.; Prinz, J.-H.; Plattner, N. J. Chem. Phys. 2013, 139, 184114.

(49) Noé, F.; Nüske, F. Multiscale Model. Simul. 2013, 11, 635–655.

(50) Nüske, F.; Keller, B. G.; Pérez-Hernández, G.; Mey, A. S. J. S.; Noé, F. J. Chem. Theory Comput. 2014, 10, 1739–1752.

(51) Wu, H.; Noé, F. arXiv:1707.04659 2017,

(52) Olsson, S.; Wu, H.; Paul, F.; Clementi, C.; Noé, F. Proceedings of the National Academy of Sciences 2017, 114, 8265–8270.

(53) Noe, F.; Doose, S.; Daidone, I.; Lollmann, M.; Sauer, M.; Chodera, J. D.; Smith, J. C. Proc. Natl. Acad. Sci. U.S.A. 2011, 108, 4822–4827.

(54) Olsson, S.; Noé, F. J. Am. Chem. Soc. 2017, 139, 200–210.

(55) Mardt, A.; Pasquali, L.; Wu, H.; Noé, F. Nature Communications 2018, 9.

(56) Wu, H.; Mardt, A.; Pasquali, L.; Noe, F. Deep Generative Markov State Models. arXiv:1805.07601, 2018.

(57) Goodfellow, I.; Bengio, Y.; Courville, A. Deep Learning; MIT Press, 2016; http://www.deeplearningbook.org.

